# Functional connectivity networks with and without global signal correction

**DOI:** 10.1101/000968

**Authors:** Satoru Hayasaka

**Author notes:** Correspondence: Satoru Hayasaka, Ph.D., Department of Biostatistical Sciences, Wake Forest School of Medicine, Medical Center Blvd., Winston-Salem, NC, 27157, USA.

## Abstract

In functional connectivity analyses in BOLD (blood oxygenation level dependent) fMRI data, there is an ongoing debate on whether to correct global signals in fMRI time series data. Although the discussion has been ongoing in the fMRI community since the early days of fMRI data analyses, this subject has gained renewed attention in recent years due to the surging popularity of functional connectivity analyses, in particular graph theory-based network analyses. However, the impact of correcting (or not correcting) for global signals has not been systematically characterized in the context of network analyses. Thus, in this work, I examined the effect of global signal correction on an fMRI network analysis. In particular, voxel-based resting-state fMRI networks were constructed with and without global signal correction. The resulting functional connectivity networks were compared. Without global signal correction, the distributions of the correlation coefficients were positively biased. I also found that, without global signal correction, nodes along the interhemisphic fissure were highly connected whereas some nodes and subgraphs around white-matter tracts became disconnected from the rest of the network. These results from this study show differences between the networks with or without global signal correction.

## 1. Introduction

Since the early days of fMRI, neuroimaging researchers have documented highly correlated time courses in distinct brain areas even when a subject is not engaged in a cognitive task. For example, Biswal et al. described strong correlation between the left and right motor cortices while the subjects were at rest (Biswal et al., 1995). Another well-documented example is a collection of brain areas, known as the default mode network (DMN), that exhibit similar time courses when subjects are at rest (Raichle and Snyder, 2007). Brain areas following a highly correlated time course despite the lack of external stimulus or cognitive engagement are often referred as *functionally connected*. Conversely *functional connectivity* between distinct brain areas can be assessed by examining the temporal correlation or coherence between the recordings from those areas. While early connectivity studies focused on functional connectivity to/from a particular seed region in the brain (for example, (Greicius et al., 2003; Fox et al., 2005; Fox et al., 2006)), in recent years, functional connectivity among different brain areas is often examined in the form of functional connectivity networks (Eguiluz et al., 2005; Salvador et al., 2005; Achard et al., 2006). Such a brain network can be constructed by examining functional connectivity originating from each distinct brain area, and organizing such connections from all the brain areas in the form of a network, with each node representing a brain area and each edge representing functional connectivity between two nodes (or brain areas) (Bullmore et al., 2009; Bullmore and Sporns, 2009; Rubinov and Sporns, 2010).

When constructing a functional brain network, it is important to process fMRI data in a way that the resulting network does not include erroneous functional connectivity resulting from confounding biases or signals not necessarily of a neurological origin. Thus, in order to construct a network, it is a common practice to pre-process fMRI data before assessing functional connectivity. In particular, a band-pass filter is applied to focus on low frequency BOLD fluctuations (Cordes et al., 2001; Fox et al., 2005; Van Dijk et al., 2010). In addition, rigid-body transformation parameters, generated during motion correction and alignment, are regressed out from fMRI time series data to lessen the impact of motion in the connectivity analysis (Fox et al., 2005). Physiologically confounding noises also need to be corrected. This is often carried out by regressing out the average time courses from the ventricles, white matter, and/or the whole-brain (Fox et al., 2005), often referred as global signals.

Among the pre-processing steps described above, regressing out global signals is somewhat controversial. The controversy stems from an argument that regressing out the average whole-brain signal inherently induces negative correlation, or anti-correlation (Murphy et al., 2009). There have been a number of studies supporting or refuting the need for global signal regression in connectivity analyses (Chang and Glover, 2009; Fox et al., 2009; Weissenbacher et al., 2009; Van Dijk et al., 2010; Anderson et al., 2011; Carbonell et al., 2011; Chai et al., 2012; He and Liu, 2012). Interestingly, some of these studies found that the distribution of correlation coefficients is positively biased (Fox et al., 2009; Murphy et al., 2009; Chai et al., 2012). Moreover, Fox et al. (2009) found that extensive brain areas are positively correlated with the whole-brain signal; this may explain the positive bias in the correlation coefficients since a large number of brain areas exhibit correlation to the same signal.

It should be noted that most of these studies described above are seed-based functional connectivity studies, in which correlation coefficients are calculated between the time series from a particular seed region and each individual voxel in the brain. On the other hand, in graph-theory-based network analyses, functional connectivity networks are constructed by calculating correlation coefficients in all possible pairs of brain areas or voxels. Thus it is still unclear how correcting for the global signal affects the resulting functional connectivity networks. During construction of functional connectivity networks, emphasis is often placed on highly positive correlations rather than negative correlations or anti-correlations. Moreover, investigators routinely select a certain proportion of high correlation coefficients to define edges in their networks; thus a positive bias in the distribution of correlation coefficients may not impact the network structure. Therefore, in this report, I investigate the impact of (not) regressing out global signals in functional connectivity networks. In particular, I constructed networks with and without global signal correction using the resting-state fMRI data from the same set of subjects. Then I examined how the network organization differed between these networks. Namely, I focused on the distribution of correlation coefficients, the locations of high degree nodes – or hubs, and the modular organization in voxel-based functional connectivity networks.

## 2. Materials and methods

### 2.1 fMRI data

I used the same dataset as the study described in Hayasaka & Laurienti (2010). I used this data set since it has been extensively studied and characterized in my previous work (Hayasaka and Laurienti, 2010). This data set consisted of fMRI time series data from 10 normal subjects (5 females, average age 27.7 years old, SD = 4.7). The fMRI data were acquired while the subjects were resting using a gradient echo echo-planar imaging (EPI) protocol with TR/TE = 2500/40 ms on a 1.5 T GE MRI scanner with a birdcage head coil (GE Medical Systems, Milwaukee, WI). Other acquisition parameters included: 24 cm field of view, and 64 × 64 acquisition matrix. The time series data included 120 images acquired over 5 minutes. The acquired images were corrected for slice timing and motion, and subsequently were realigned. Then the images were spatially normalized to the MNI (Montreal Neurological Institute) space and re-sliced to 4 × 4 × 5 mm voxel size using an in-house processing script based on the SPM package (Wellcome Trust Centre for Neuroimaging, London, UK). The resulting fMRI time series data were band-pass filtered (0.009-.08 Hz) to attenuate respiratory and other physiological noises. These processing steps are widely used in fMRI functional connectivity studies (Fox et al., 2005; van den Heuvel et al., 2008). More details on my data pre-processing steps can be found elsewhere (Hayasaka and Laurienti, 2010; Joyce et al., 2010).

### 2.2 Global signal regression

I considered four different methods of global signal correction. Although there are many possible ways of correcting global signals, examining a large number of such methods may be beyond the scope of this work. Thus I focused on the methods that have been widely used in the literature examining the impact of global signal correction (Chang and Glover, 2009; Fox et al., 2009; Murphy et al., 2009; Weissenbacher et al., 2009; Van Dijk et al., 2010; Anderson et al., 2011; Chai et al., 2012; He and Liu, 2012; Hallquist et al., 2013). Mean time courses from the entire brain (the average of voxel values within the brain parenchyma mask including gray and white matter), the deep white matter (average time course in an 8 mm radius sphere within the anterior portion of the right centrum semiovale composed entirely of white matter), and the ventricles (average of time courses within the ventricle mask) were extracted and used in global signal correction as described below. In the first method, 6 rigid-body transformation parameters, generated during the realignment (note: NOT normalization) step, were regressed out from the fMRI time series data (Fox et al., 2009). This method was referred as the no correction method (NoCorr), since no global signals, besides the motion parameters, were regressed out from the data. This method demonstrated a situation in which global signal correction is completely omitted. In the second method, in addition to the motion parameters as described above, the average time course from the deep white matter and the ventricles were regressed out, but not the average whole-brain signal (Chang and Glover, 2009; Fox et al., 2009; Weissenbacher et al., 2009; Anderson et al., 2011; He and Liu, 2012). This method was referred as the no whole-brain signal method (NoWB). In the third method, only the whole-brain signal was regressed out in addition to the motion parameters (Murphy et al., 2009; Van Dijk et al., 2010; Anderson et al., 2011; He and Liu, 2012). This method was referred as the whole-brain only method (WBonly). Finally, in the fourth method referred as the full method (Full), the motion parameters as well as the average signals from the white matter, ventricles, and whole-brain were regressed out (Fox et al., 2009; Van Dijk et al., 2010; Chai et al., 2012; Hallquist et al., 2013). The Full networks served as the baseline in this study, characterizing differences in the network organization when one or more global signal variables are omitted. Figure 1 describes the overview of the different methods. It was noted by one of the reviewers that regression after filtering has been criticized by some studies (Hallquist et al., 2013; Saad et al., 2013).

**Figure 1:**
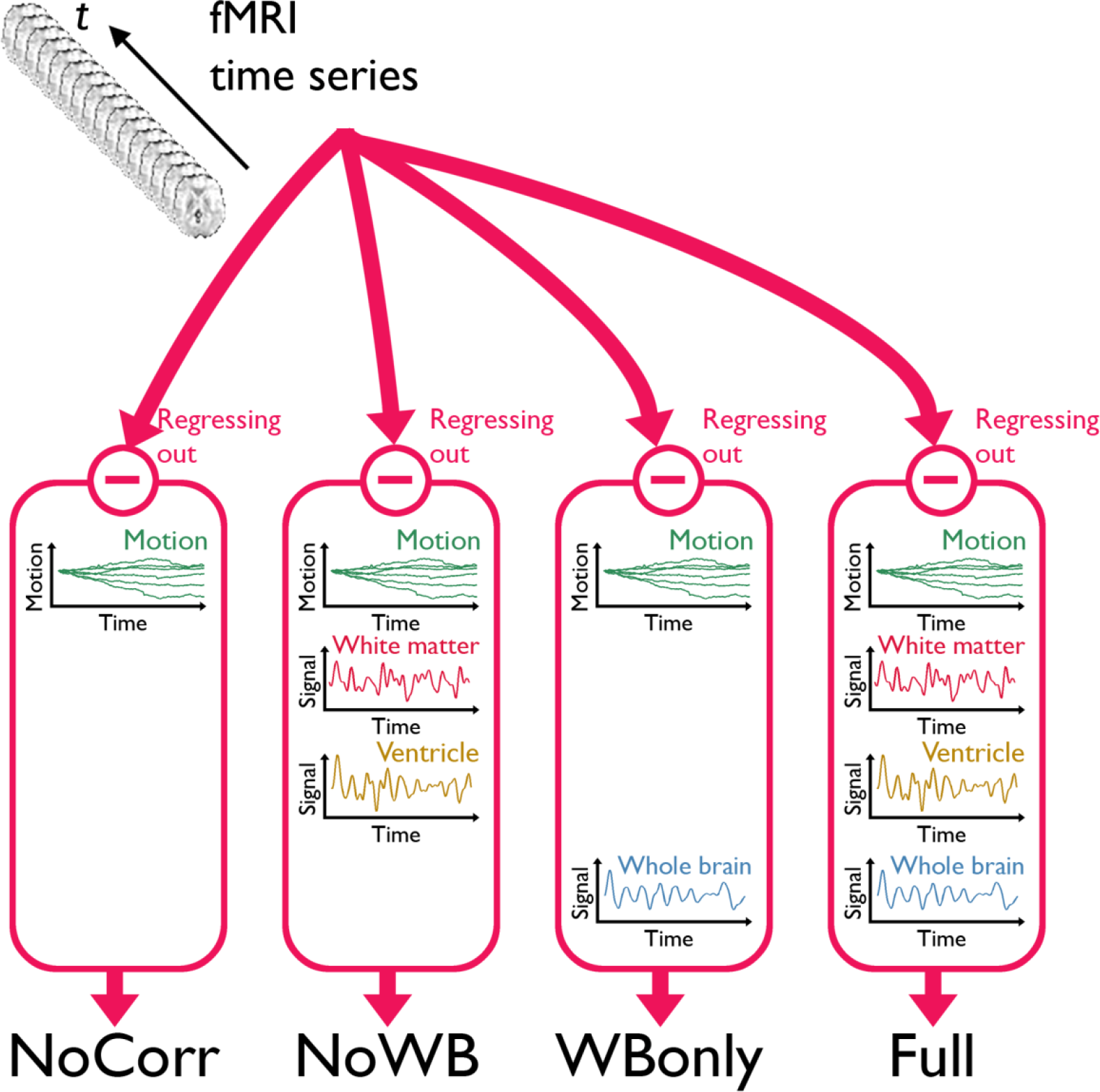
A schematic of the three global signal correction methods. Motion corrected and band-pass filtered fMRI time series data from each subject were processed in four different methods to correct for global signals. In the first method, only the 6 parameters associated with a rigid-body transformation were regressed out from the fMRI time series (NoCorr). In the second method, in addition to the motion parameters, the mean signals from the deep white matter and the ventricles were also regressed out (NoWB). In the third method, only the whole-brain signal is regressed out in addition to the motion parameters (WBonly). In the fourth method, the motion parameters as well as the mean signals from the white matter, the ventricles, and the whole-brain were regressed out (Full).

### 2.3 Network construction

Processed in one of the four methods described above, the fMRI time series data from each subject were then used to construct a functional brain network, with each node representing a voxel and each edge representing a strong linear correlation between two voxel time courses. To ensure all the networks from all the subjects have the same set of nodes, a binary mask image was generated comprising 15,996 voxels within the AAL (automated anatomical labeling) atlas (Tzourio-Mazoyer et al., 2002). Among these voxels within the mask, a cross-correlation matrix was calculated, with each element being the correlation coefficient between two voxel time courses. The resulting correlation matrix consisted of 255,856,020 correlation coefficients (excluding the main diagonal elements, which are 1).

I then examined the distribution of the correlation coefficients in the correlation matrix. The exact marginal distribution of each correlation coefficient *r* is

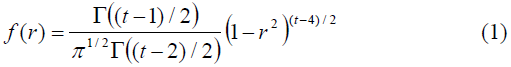

where *t* is the number of time points (Johnson et al., 1995; Cao and Worsley, 1999). However, correlation coefficients in the correlation matrix are not independent. Rather, collectively they represent a 6-dimensional “*connexel*” field (Worsley et al., 1998; Cao and Worsley, 1999). Consequently the collective distribution of all the correlation coefficients in this correlation matrix does not follow (1). Nevertheless, since the marginal distribution (1) is centered around 0 and symmetric, the histogram of the correlation coefficients should be centered at 0 and symmetric. Any deviation from mean = 0 can be an indication of a systematic bias in the correlation matrix. Or, if there is a true global signal present in all the voxels, that also biases the distribution of correlation coefficients. Let W_1_ = Y_1_ + G and W_2_ = Y_2_ + G be two voxel time courses, where Y_1_ and Y_2_ indicate intrinsic time courses in both voxels and G is the global signal present in both W_1_ and W_2_. If the global signal G is uncorrelated with neither Y_1_ nor Y_2_ (i.e., Cov(Y_1_,G) = 0 and Cov(Y_2_,G) = 0), then the covariance between the two voxel time courses W_1_ and W_2_ is

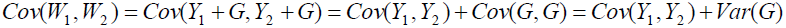

The variance of W_1_ and W_2_ are Var(W_1_) = Var(Y_1_) + Var(G) and Var(W_2_) = Var(Y_2_) + Var(G), respectively. Thus, even if Y_1_ and Y_2_ are uncorrelated (i.e., Cov(Y_1_,Y_2_) = 0), the correlation coefficient between W_1_ and W_2_

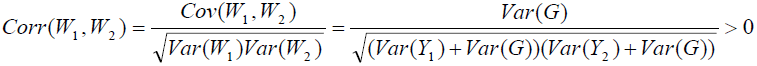

is always positive since Var(G) is always positive. Because of this, the distribution of correlation coefficients in this case no longer follows (1) but follows a non-central form

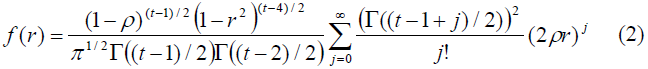

where ρ = Corr(W_1_,W_2_). This distribution is no longer symmetric around 0. In the literature on functional connectivity, there have been some reports that the distribution of correlation coefficients is positively biased when global signals are not corrected (Fox et al., 2009; Murphy et al., 2009; Chai et al., 2012). Thus, to examine whether there is such a systematic bias, I generated a histogram of the correlation coefficients for each method (NoCorr, NoWB, WBonly, or Full) for each subject. The means from the correlation coefficient distribution were compared across different correction methods by paired two-sample t-tests.

The correlation matrix from each subject and each correction method was then used to construct a functional connectivity network. In particular, the correlation matrix was thresholded to generate a binary adjacency matrix with 1 indicating the presence and 0 indicating the absence of an edge between two nodes, with each edge representing a strong positive correlation. I chose a positive threshold in a way to control the number of nodes N and the average node degree K in the resulting network. In particular, I selected a correlation threshold such that the ratio S = log(N)/log(K) is the same across subjects. I chose S = 3.0 since it has been shown to capture the network characteristics effectively (Hayasaka and Laurienti, 2010) and the resulting edge density is comparable to that of a self-organized network of a similar size (Laurienti et al., 2011). I examined the results with different values of S ranging between 2.5 and 3.5, and the results were similar across S values in comparisons of network characteristics across the methods (results not shown). Thus, throughout this paper, only the results for the networks with S = 3.0 are shown.

Once the network was generated, various network characteristics were compared. This includes whole-network metrics such as clustering coefficients C and path length L (Watts and Strogatz, 1998; Stam and Reijneveld, 2007). While C represents the probability that a node’s neighbors are also neighbors to each other, L is the average of shortest distances between any two nodes in a network, in terms of the number of edges separating them or the geodesic distance. These metrics were compared across different methods by paired two-sample tests. Moreover, I examined the consistency of high degree nodes, or hubs, across subjects. This was done by examining the spatial overlap of top 20% highest degree nodes across subjects (Hayasaka and Laurienti, 2010). The resulting overlap images were compared across different correction methods. If global signal correction does not influence the overall structure of the network, then the overlap maps should appear similar across different correction methods. On the other hands, if systematic biases are introduced by global signal correction, or by the lack thereof, then the overlap maps may appear different across the correction methods.

### 2.4 Modular organization

In a network, some groups of nodes may have a large number of connections among themselves compared to connections between such groups. These highly interconnected sets of nodes are often referred as modules. If a network has a modular structure, then its nodes can be grouped into a number of modules, with each node belonging to a single module. The human brain networks have been shown to have modular organization (He et al., 2009; Meunier et al., 2009; Power et al., 2011; Rubinov and Sporns, 2011). Despite the difference in the number of nodes in these previous studies, the number of modules is similar and the modular parcellation is comparable (Moussa et al., 2012). Thus, I hypothesize that, if a lack of global signal correction alters the macro-scale organization of a functional brain network, such altered organization may manifest as changes in the modular organization.

To investigate the modular organization, I applied an algorithm called Qcut (Ruan and Zhang, 2008). Qcut is an iterative algorithm to find a near optimal modular parcellation of a network, by maximizing modularity Q, a metric that quantifies how parcellated a network is relative to a random network of a comparable size. Q is zero if the network exhibits no community structure, whereas a large Q is a strong indicator of community structure in a network (Clauset et al., 2004). The upper limit of Q is 1. For each fMRI network, before running Qcut, I identified sub-networks that were isolated from the largest connected network component (or the giant component), and grouped such nodes into a “*junk*” module. Then the giant component was analyzed by Qcut, resulting in a modular parcellation. The resulting Q was compared across different correction methods, and the consistency of some modules was examined.

## 3. Results

### 3.1 Correlation coefficient distribution

Figure 2 shows distributions of correlation coefficients for all the subjects under different correction methods. While the distribution was centered at 0 for all the subjects for the Full and WBonly methods, the distribution was positively skewed in some subjects for the NoWB and NoCorr methods. Between NoWB and NoCorr, the distribution appeared more skewed for NoCorr. This was confirmed by the mean of these distributions. The mean (SD) of the mean correlation coefficient across subjects was 0.00006 (0.0004) under the Full method, 0.00006 (0.0004) under the WBonly method, 0.050 (0.035) under the NoWB method, and 0.086 (0.056) under the NoCorr method. I compared the mean correlation coefficient between different methods by paired two sample t-tests (since the networks originate from the same set of subjects). I found a significant difference between the Full and NoWB methods (p = 0.001), as well as between the Full and NoCorr methods (p<0.001). However, no significant difference was found between the Full and WBonly methods (p = 0.70). Significant differences were also found between WBonly and NoWB methods (p = 0.001) as well as between the WBonly and NoCorr methods (p<0.001). These results indicate that the correlation matrix may be systematically biased when the whole-brain signal is not regressed out. These results are consistent with previous reports on seed-based connectivity studies (Fox et al., 2009; Murphy et al., 2009; Chai et al., 2012).

**Figure 2:**
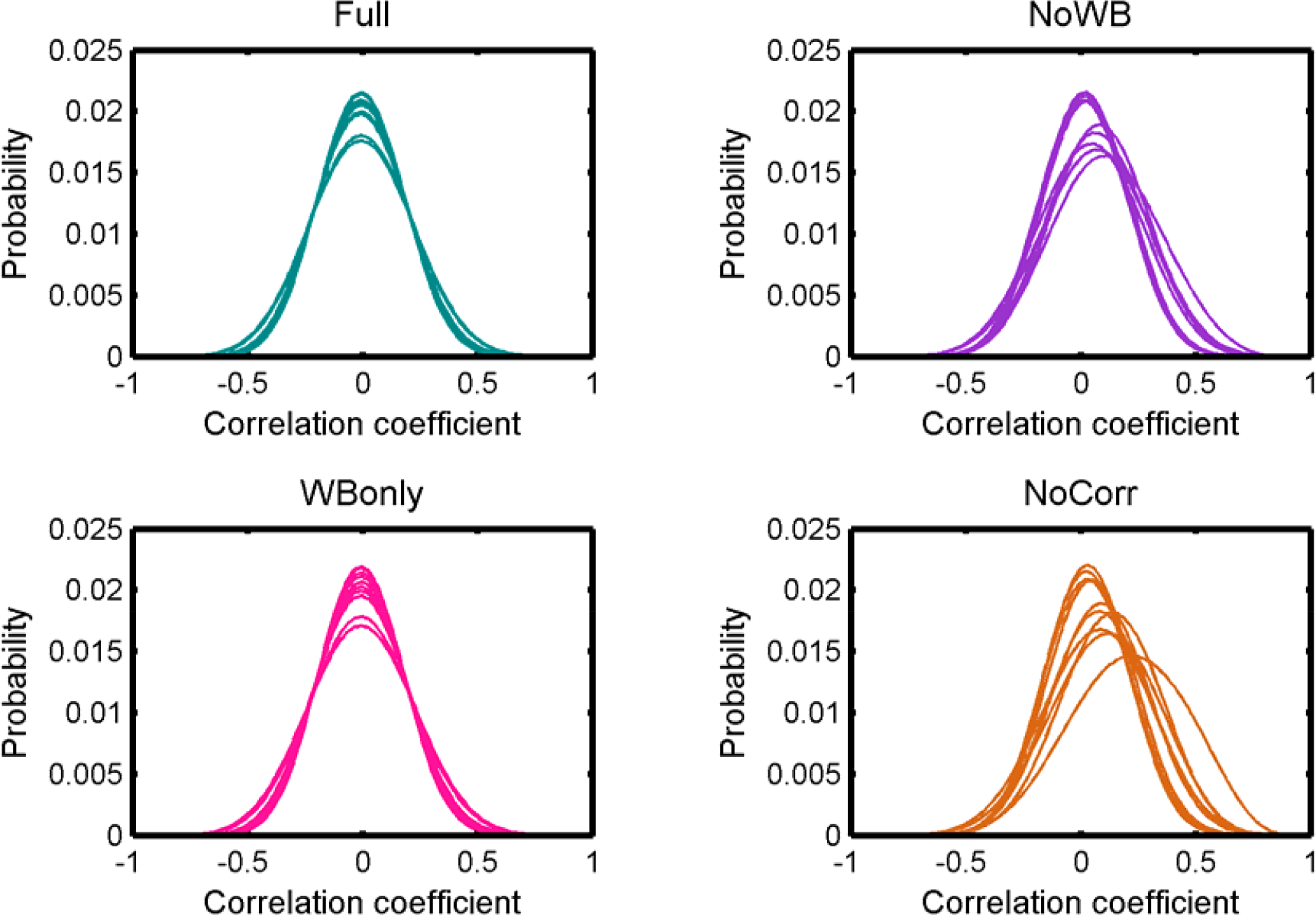
Distributions of correlation coefficients. Distributions of correlation coefficients from different subjects’ networks under different global signal correction methods. While the distributions are centered at zero for the Full and WBonly methods, the distributions are positively skewed for some subjects for the NoWB and NoCorr methods.

### 3.2 Network metrics

Table 1 shows the average C and L for the four different methods. While clustering coefficient C appeared similar across different correction methods, path length L was somewhat larger for the NoWB and NoCorr networks, in comparison to the Full and WBonly networks. Paired two-sample t-tests revealed that the path lengths were marginally larger for the NoWB, WBonly, and NoCorr networks when compared to that of the Full networks (p = 0.09, p = 0.03, and p = 0.05, respectively). This may be because the NoWB, WBonly, and NoCorr networks fragmented more than the Full networks. In fact, the size of the largest connected network component Nc, or the size of the giant component, was smaller in the NoWB, WBonly, and NoCorr networks compared to the Full networks (paired t-test p = 0.03, p = 0.02, and p = 0.006, respectively) (see Table 1). Since my method of path length calculation was based on the reciprocal mean of the geodesic distance between nodes (Latora and Marchiori, 2001; Hayasaka and Laurienti, 2010), disconnected network components were accounted as increased path length. Furthermore, the proportion of connected nodes (i.e., nodes with at least one connection) was much lower in the NoWB and NoCorr networks compared to the Full networks (paired t-test p = 0.04 and p = 0.01, respectively) (see Table 1). However, the proportion of connected nodes was only marginally smaller in the WBonly networks compared to the Full networks (paired t-test p = 0.08). It should be noted that the difference in the path length L as described above cannot be simply attributed to the differences in the distribution of correlation coefficients. This is because a distribution of correlation coefficients does not describe the network structure or topology, as it lacks information on how nodes are connected to each other.

**Table 1:** Average network metrics. The average network metrics across subjects were calculated for the networks with different global signal correction methods. Clustering coefficient C and path length L, as well as the size of the giant component Nc and the proportion of nodes with at least one connection are presented.

| Correction method | C<br>Mean (SD) | L<br>Mean (SD) | Nc<br>Mean (SD) | Proportion of<br>connected nodes<br>Mean (SD) |
| --- | --- | --- | --- | --- |
| Full | 0.230 (0.039) | 5.29 (1.01) | 14897 (916) | 94% (4%) |
| NoWB | 0.223 (0.031) | 7.35 (3.90) | 13246 (2430) | 86% (13%) |
| WBonly | 0.236 (0.039) | 5.49 (1.18) | 14699 (1095) | 94% (5%) |
| NoCorr | 0.224 (0.032) | 8.72 (5.11) | 12245 (2616) | 81% (15%) |

### 3.3 Node degree distribution

Figure 3 shows the degree distributions for the networks constructed with different correction methods. In all the methods, the degree distributions seem to follow an exponentially truncated power-law distribution, as I previously reported (Hayasaka and Laurienti, 2010). However, the shape of the distributions appeared more variables in the NoWB and NoCorr networks. To confirm this, the variance of the largest node degree was compared across different correction methods by an F-test. The variability was significantly larger in the NoWB and NoCorr networks compared to the Full networks (F-test p = 0.008 and p = 0.005, respectively), or compared to the WBonly networks (F-test p = 0.01 and p = 0.008, respectively). However, no significant difference in variability was found between the Full networks and the WBonly networks (F-test p = 0.82).

**Figure 3:**
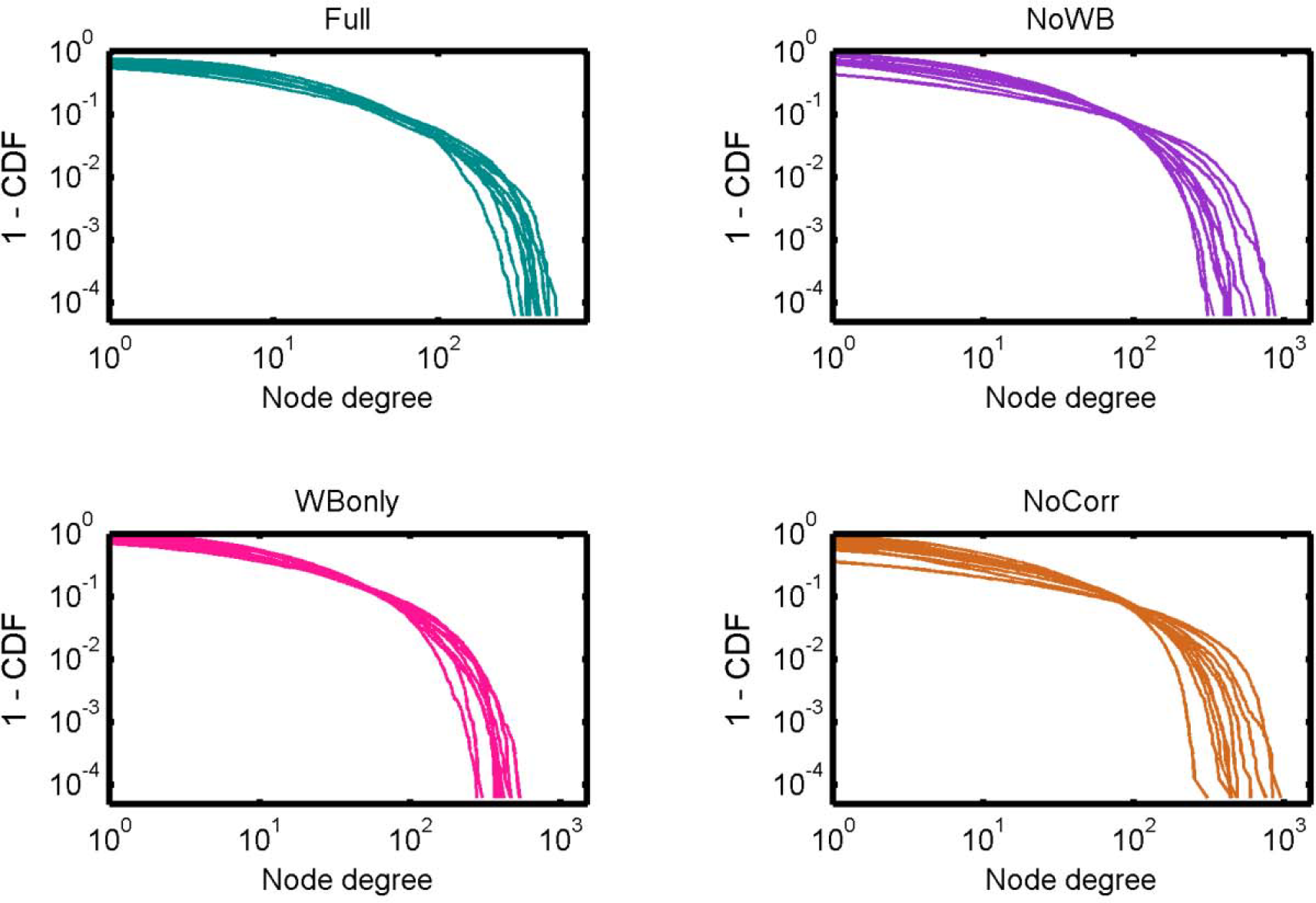
Degree distributions. Degree distributions of the Full, NoWB, WBonly and NoCorr networks are shown. Although all the distributions seem to follow an exponentially truncated power-law distribution, the degree distributions appear more variable across subjects in the NoWB and NoCorr networks.

### 3.4 Network hubs

Next, I examined the locations of high-degree nodes, or hubs, in the networks with different correction methods. In particular, the consistency of hub locations was examined by an overlay image of top 20% highest degree nodes (see Figure 4). All the methods yielded a concentration of network hubs in the posterior cingulate cortex and the precuneus. This finding was consistent with my previous results (Hayasaka and Laurienti, 2010) as well as the other voxel-based network studies (Eguiluz et al., 2005; van den Heuvel et al., 2008; Buckner et al., 2009). However, the NoWB and NoCorr networks also showed a concentration of hub nodes near the superior edge of the interhemispheric fissure while such concentration was not observed in the Full and WBonly networks. To the best of my knowledge, this area has not been reported as the hub area of the brain in voxel-level fMRI networks. Moreover, resting-state MEG (magnetoencephalography) networks often do not exhibit concentration of hubs along the interhemispheric fissure (Bassett et al., 2006; Deuker et al., 2009; Jin et al., 2013; Rutter et al., 2013). The concentration of hub nodes in this area was more consistent and extensive in the NoCorr networks than the NoWB networks. Thus it is possible that this concentration is an artifact of not correcting for the whole-brain signal. It should also be noted that, while the Full networks showed a concentration of hub nodes in the anterior cingulate cortex, the NoWB, WBonly, and NoCorr networks did not show such a concentration in the same area.

**Figure 4:**
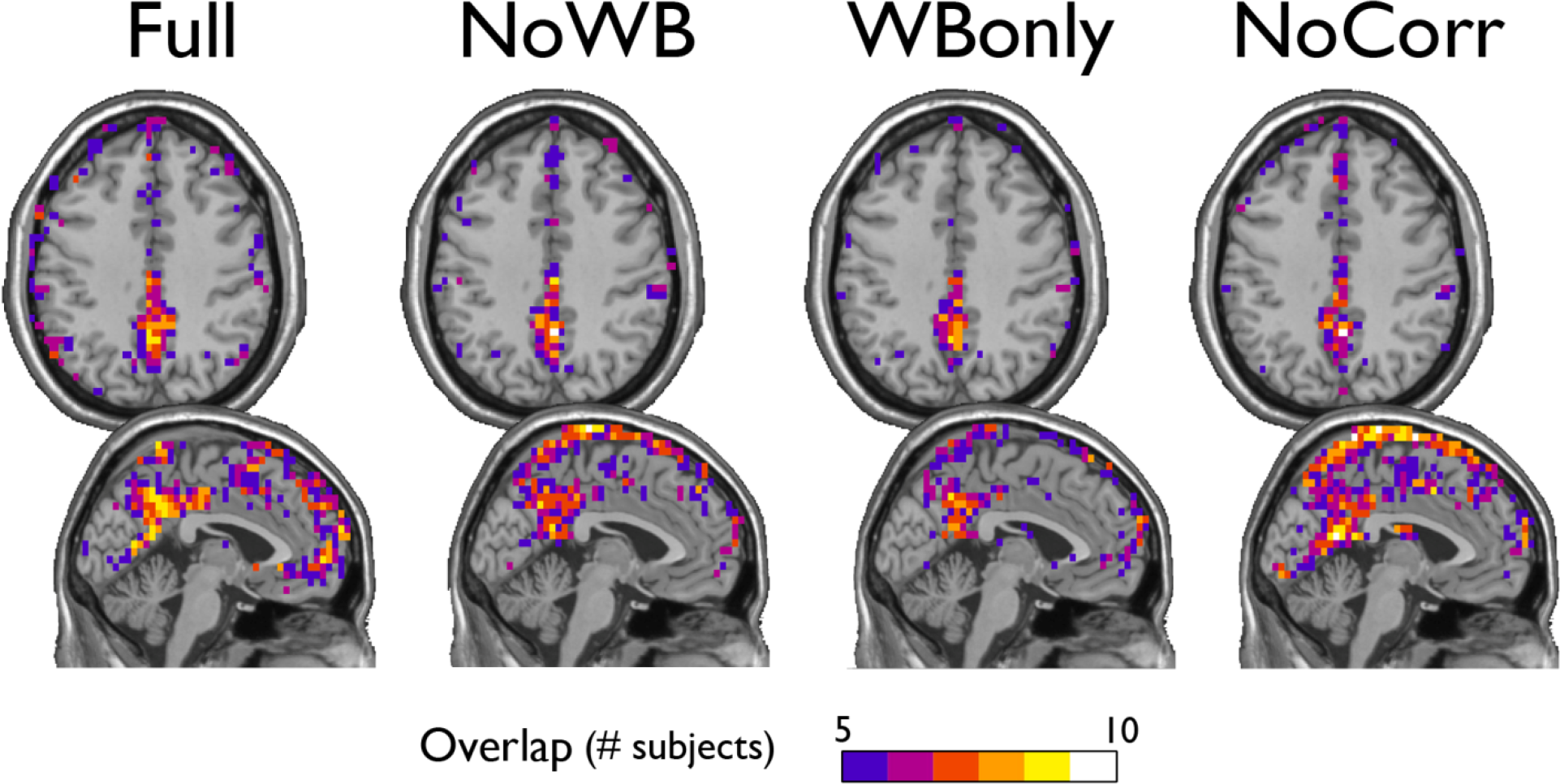
Consistency of hub nodes. The overlap of hub nodes (top 20% highest degree nodes) across subjects is shown for the networks with different correction methods. Hub nodes were consistently concentrated in the posterior cingulate cortex and the precuneus. However, the NoWB and NoCorr networks also showed a concentration of hub nodes along the superior edge of the interhemispheric fissure, while such a concentration was not observed among the Full and WBonly networks.

### 3.5 Modular organization

Table 2 shows the mean modularity Q of the networks under different correction methods, as well as the mean number of modules found in these networks. Compared to the Full networks, modularity Q did not differ significantly in the NoWB, WBonly, and NoCorr networks (paired t-test p-values, p = 0.12, p = 0.20, and p = 0.05, respectively). However, there were significantly more modules in the NoWB and NoCorr networks compared to the Full networks (paired t-test p = 0.02 and p = 0.004, respectively). Compared to the WBonly networks, the NoWB and NoCorr networks had significantly more modules as well (paired t-test p = 0.03 and p = 0.01, respectively). There was no significant difference in the number of modules between the Full and WBonly networks (p = 0.08). These results indicate that the brain network is parcellated into a larger number of communities when the whole-brain signal is not corrected.

**Table 2:**
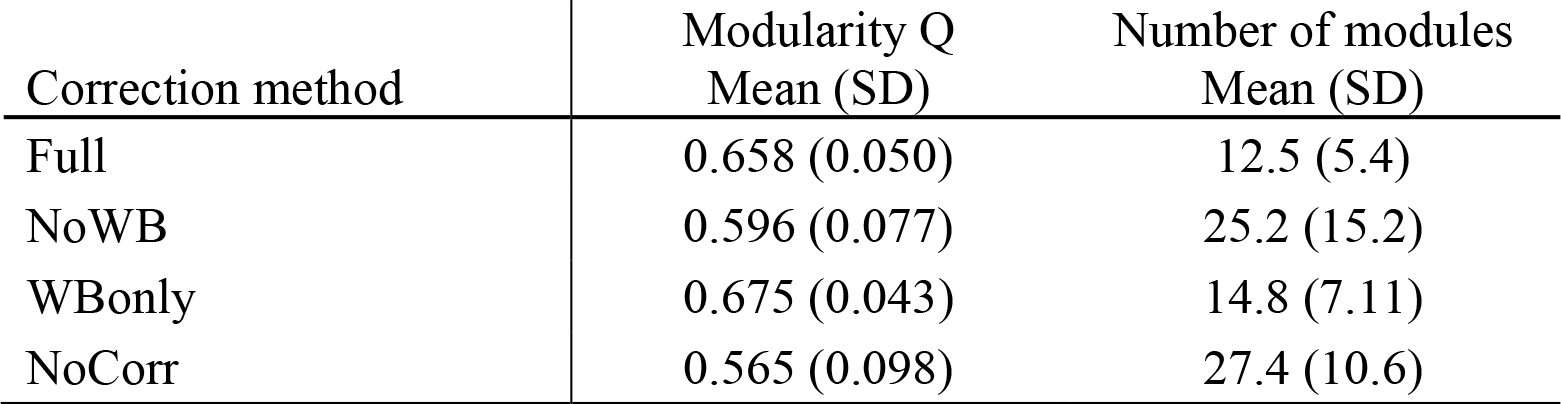
Modularity Q and the number of modules. The mean modularity Q from the networks under different correction methods, as well as the mean number of modules are shown.

I examined the consistency of the default mode network DMN module across subjects under different correction methods. In particular, for each method, I generated an overlap image of the DMN module, identified manually as the module comprising a large portion of the posterior cingulate cortex and the precuneus, the areas known to be part of the DMN. Figure 5 shows the overlap images demonstrating the consistency of the DMN. Surprisingly, the DMN overlap images were similar across different methods. This result indicates that the difference in correction methods did not impact the DMN module.

**Figure 5:**
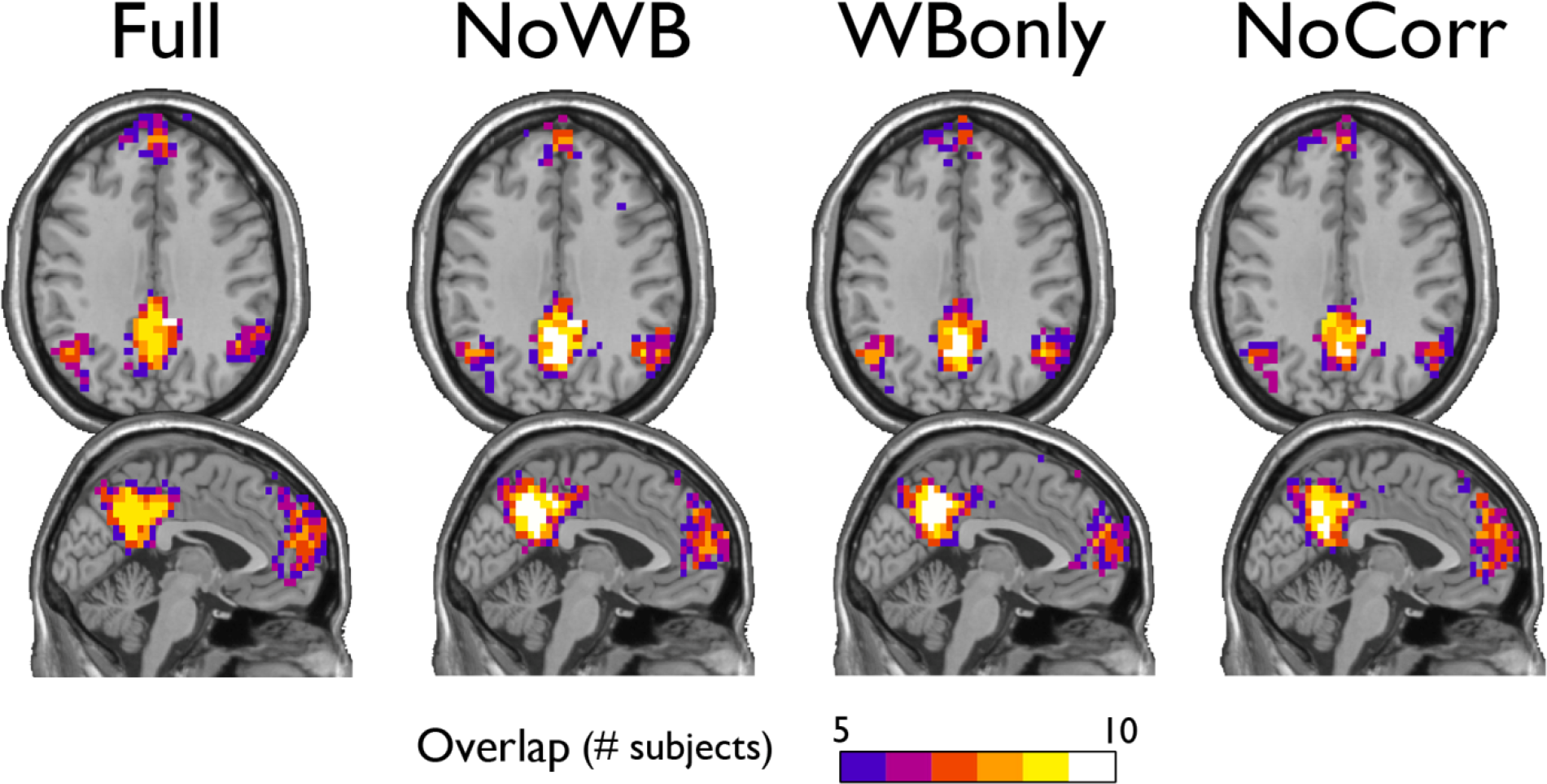
Consistency of the default mode network module. The overlap of the default mode network module across subjects is shown for the networks with different correction methods. The areas of overlap appear similar across different methods.

I also examined the consistency of the junk module, the module consisting of nodes and subgraphs disconnected from the giant component of the brain network. Figure 6 shows the overlap images of the junk module across subjects under different correction methods. While the junk module was not spatially consistent across subjects in the Full and WBonly networks, the junk module consistently included nodes around the major white matter tracts in the NoWB and NoCorr networks. It should be noted that, my network data only consisted of gray matter nodes defined by the AAL atlas. Between the NoWB and NoCorr networks, the overlap was more consistent and extensive in the NoCorr networks. To further investigate these differences, I counted the number of nodes in the junk module (i.e., isolated nodes and subgraphs) that are adjacent to major white matter tracts. Such nodes adjacent to white matter tracts were identified from the gray matter voxels constituting a brain network (Figure 7, left). Among these voxels, ones with white matter probability greater than 40% were identified using the white matter probability map from the SPM package; the resulting mask included nodes that were adjacent to white matter tracts, as it can be seen in Figure 7, right. The average numbers of junk module nodes within this mask for different correction methods are shown in Table 3. Compared to the Full networks, there were more junk module nodes (i.e., isolated nodes and subgraphs) adjacent to white matter tracts in the NoWB, WBonly and NoCorr networks (paired t-test p = 0.02, p = 0.02, and p = 0.002, respectively). Compared to the WBonly networks, the NoWB and NoCorr networks had significantly more junk module nodes adjacent to white matter tracts (paired t-test p = 0.02, and p = 0.003, respectively). These results indicated that, without regressing out the whole-brain signal, some nodes may be systematically disconnected from the rest of the network, especially around white matter tracts.

**Figure 6:**
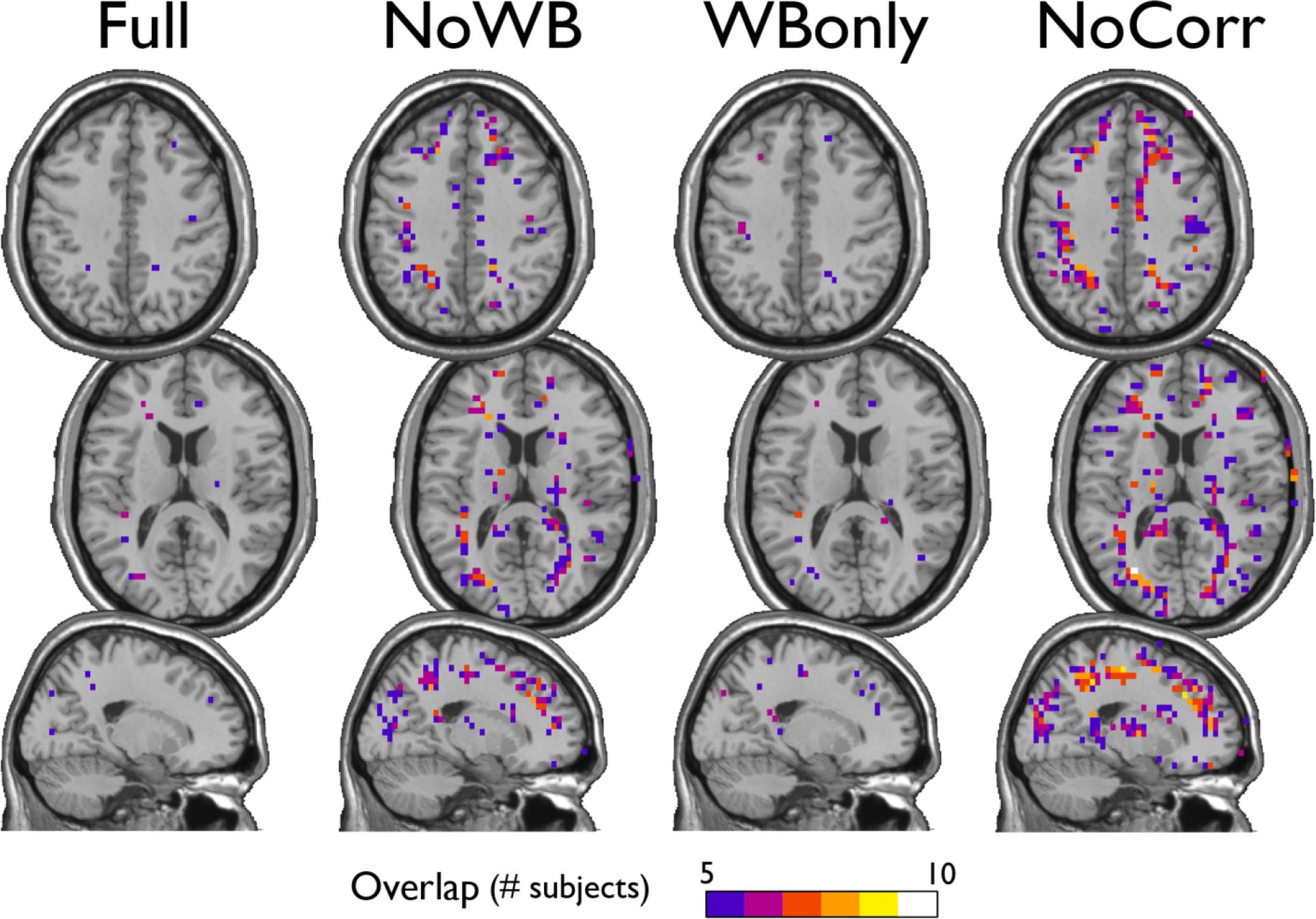
Consistency of the junk module. The junk module in each network consists of nodes and subgraphs that are disconnected from the giant component. Under each global signal correction method, the consistency of such junk modules across subjects was examined by generating an overlap image. While the junk module showed only isolated signs of consistency in the Full and WBonly networks, the junk module consistently included nodes around the major white matter tracts in the NoWB and NoCorr networks. Between the NoWB and NoCorr networks, the overlap was more consistent and extensive in the NoCorr networks.

**Figure 7:**
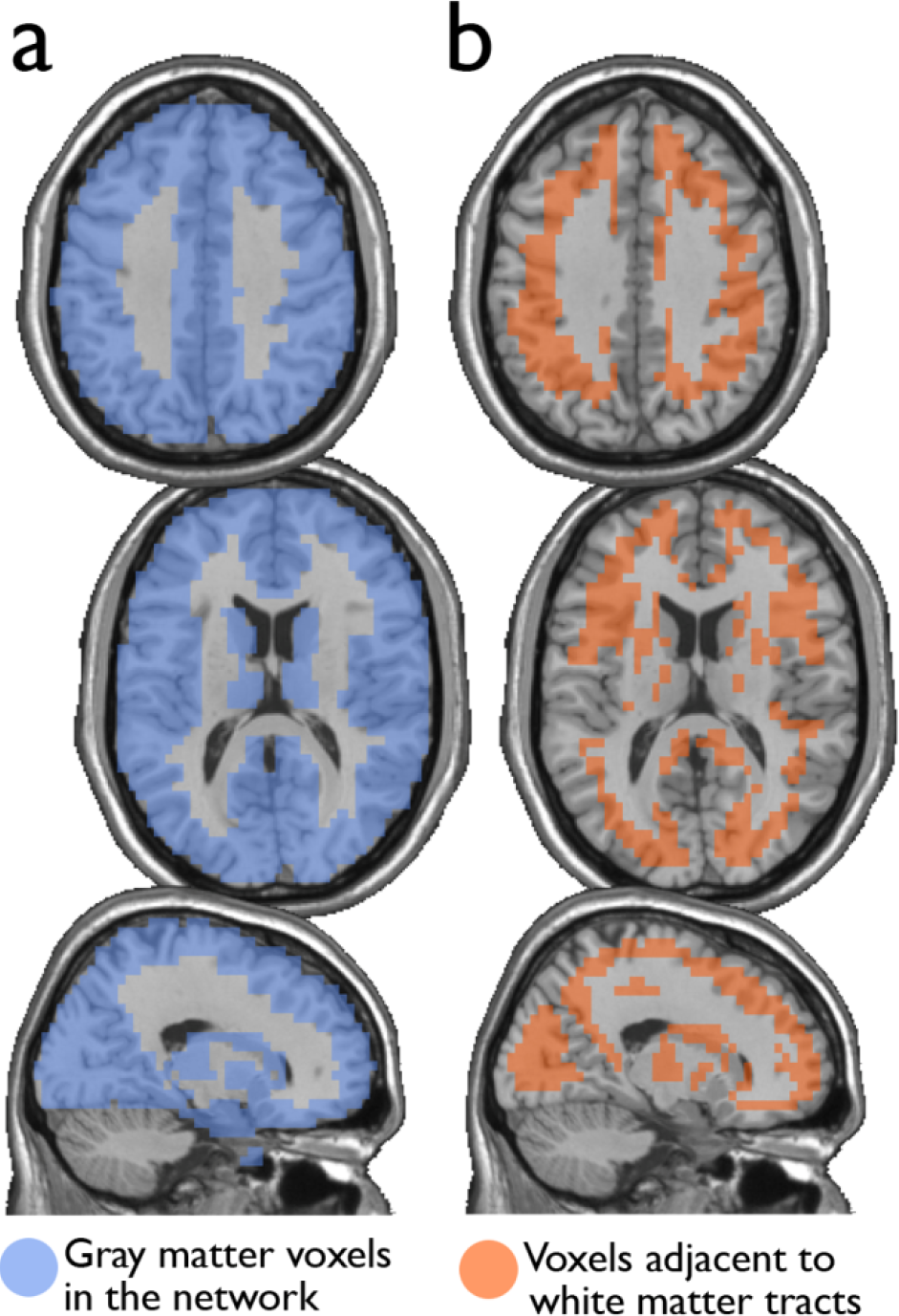
Nodes surrounding white matter tracts. Among the gray matter voxels included as part of a brain network (left),voxels adjacent to the major white matter tracts were identified (right). These voxels were identified from a white matter mask image from the SPM package, with at least 40% white matter probability.

**Table 3:** The number of junk module nodes adjacent to white matter tracts. The number of nodes within the junk modules which are within the white matter adjacency mask (Figure 7, right) is listed for different correction methods.

| Correction method | Number of Junk Module Nodes adjacent to<br>White Matter Tracts<br>Mean (SD) |
| --- | --- |
| Full | 384 (324) |
| NoWB | 865 (684) |
| WBoonly | 448 (377) |
| NoCorr | 1146 (675) |

Next, I examined the accuracy of the gray matter mask used in this study (i.e., voxels belonging to areas identified by the AAL atlas (Tzourio-Mazoyer et al., 2002)). This was done by first eliminating the nodes adjacent to major white matter tracts (Figure 7, right) from the whole-brain networks, and then by comparing path length L of the resulting networks to that of the whole-brain networks, as suggested by one of the reviewers. As mentioned above, the whole-brain network consisted of 15,996 nodes, whereas the networks without nodes adjacent to white matter tracts consisted of 12,660 nodes. In other words, the network size was reduced by 20%. The path lengths L for the network with and without the nodes adjacent to white matter tracts are shown in Table 4, along with the p-values from a paired t-test comparing them. While the path length L was significantly shorter without nodes adjacent to white matter tracts in the NoCorr networks (p = 0.038), no significant difference was found in the other correction methods. The difference may simply be a result of a reduced network size, or there may be a systematic connectivity difference in nodes adjacent to white matter tracts.

**Table 4:** The path length L of the networks with and without nodes adjacent to major white matter tracts. The path length L was compared between the two types of networks by a paired t-test. The resulting p-values are also shown.

| Correction method | Path Length L,<br>Networks with Nodes<br>adjacent to White Matter<br>Tracts<br>Mean (SD) | Path Length L,<br>Networks without Nodes<br>adjacent to White Matter<br>Tracts<br>Mean (SD) | P-value,<br>Paired T-test |
| --- | --- | --- | --- |
| Full | 5.29 (1.01) | 5.24 (0.88) | 0.365 |
| NoWB | 7.35 (3.90) | 6.94 (3.34) | 0.053 |
| WBoonly | 5.49 (1.18) | 5.40 (1.01) | 0.175 |
| NoCorr | 8.72 (5.11) | 8.05 (4.27) | 0.038 |

## 4. Discussion

I have constructed voxel-based functional brain connectivity networks from the same set of resting-state fMRI data but with four different methods of global signal correction. I found that the correlation coefficients were positively biased in the methods without the whole-brain signal correction. The bias was stronger if no global signal was corrected at all. I also found that, without correcting the whole-brain signal, the resulting networks may include a large number of isolated nodes and subgraphs disconnected from the giant component. This resulted in increased path length L, with a stronger effect on the NoCorr networks than the NoWB networks. While high degree nodes, or hub nodes, were consistently observed in the posterior cingulate cortex as previously reported regardless of the correction method, the networks without whole-brain signal correction exhibited consistent concentration of hub nodes along the superior portion of the interhemispheric fissure. Since this area has not been reported as the hub region in previous research, especially the ones based on neuromagnetic activities observed by MEG, it is likely that such a concentration of hub nodes may be an artifact resulting from a lack of whole-brain signal correction. I also examined the modular organization of the networks with different correction methods, and found that the networks without whole-brain signal correction were parcellated into a larger number of modules. Despite the difference in global signal correction, the DMN module was observed consistently across subjects. I also found that, in the networks without full global signal correction (whole-brain signal in particular), nodes near the major white matter tracts were systematically disconnected from the rest of the network. This was particularly apparent in the NoCorr networks.

As described above, there are some different characteristics between the networks with and without whole-brain signal correction. One possible explanation for such differences is the highly connected area along the superior edge of the interhemispheric fissure in the NoWB and NoCorr networks, in comparison to the Full or WBonly networks. Since the number of edges in a network is indirectly controlled by the way the correlation matrix is thresholded (see 2.3), an abundance of edges in one area of the network can result in reduced edges in other areas of the network. From Figure 4, I can infer that extra edges were allocated near the interhemispheric fissure in the NoWB and NoCorr networks, and these extra edges would deprive connections in other areas of the brain. This resulted in a larger number of disconnected nodes and subgraphs in the NoWB and NoCorr networks compared to the Full or WBonly networks. Such disconnected components concentrate around the white matter tracts, as it can be seen in Figure 6. These alterations appeared more pronounced in the NoCorr networks than the NoWB networks. This may be because the NoWB networks are corrected by global signals to a certain degree, while the NoCorr networks are not adjusted by any global signals at all. Such a systematic fragmentation around white matter tracts can be observed even in the WBonly networks, when compared to the Full networks. Despite the alterations in the number of connections as described above, the modular organization of the NoWB and NoCorr networks was not completely altered. In fact, possibly because of the modular nature of the brain networks, the DMN module in the NoWB and NoCorr networks was surprisingly similar to that of the Full or WBonly networks (see Figure 5).

In this study, I focused on alterations in various network characteristics when resting-state fMRI data were not corrected for global signals, compared to that of the networks constructed with a global signal correction method regressing out whole-brain, white matter, and ventricle signals (Fox et al., 2006; Fox et al., 2009). However, global signal correction method used for the Full networks is far from perfect. This correction method entails simply regressing out mean signals from the fMRI time series, which is more simplistic than methods using the physiological signals recorded during the fMRI data acquisition (Chang and Glover, 2009). Rather than regressing out the global signals, perhaps a more sophisticated approach, such as principal component analysis (PCA) or independent component analysis (ICA), may be effective in extracting neurologically relevant data from physiological noises (Chai et al., 2012). Although these shortcomings exist, regression-based methods are easy to implement as a part of cross-correlation calculation, since it only involves regressing out a number of nuisance covariates from the fMRI time series. These global covariates can be calculated from the fMRI data itself; thus this would be ideal for re-analyzing fMRI data acquired without the accompanying physiological recording. Thus this type of global signal correction method would be amenable to various types of existing fMRI data, even those that are publicly available for downloading.

It should be noted that there are infinitely many ways of correcting for global signals, and the four methods presented in this work simply represent popular methods used among neuroimaging researchers. It is possible that there are other correction methods suitable for constructing functional connectivity network. However, the goal of this paper is to evaluate existing methods; my intention is not to develop better correction methods. With increased interests in this field in recent years, it is possible that some brain network researchers will develop methods more suitable than the ones examined in this work.

One limitation in this work is the lack of ground truth in evaluating different correction methods. This is due to computational challenges arising from generating a gold standard with thousands of time series (each corresponding to a voxel time course) with a small number of known correlations among them representing the “true” connectivity in an adjacency matrix. This is a very difficult mathematical problem, as if there were thousands of simulated regions in the simulation described in Saad et al. (2012) and each region’s connectivity would have to exactly match the ground truth adjacency matrix.

I also would like to emphasize that this study does not answer whether or not there is a genuine “global signal” that is present throughout the brain. This study only outlines the differences in network organization arising from correcting / not correcting for global signals. There are a number of papers describing the existence of such global signals and consequently discouraging the use of global signal correction (Murphy et al., 2009; Scholvinck et al., 2010; Saad et al., 2012; Hallquist et al., 2013; Saad et al., 2013). Because of the limitations listed above, I cannot conclude which correction method should be used, if used at all. So I will leave that determination up to each reader. If one suspects that there exists a true “global signal” that covers extensive cortical areas due to a brain-wide synchronized neurological processing, then a global signal regression is not necessary. However, I would like to reiterate that, without global signal correction, a concentration of hubs appears at the superior portion of the interhemispheric fissure, which cannot be detected by MEG. Moreover, nodes around white matter tracts tend to systematically disconnect from the rest of the brain network if the whole-brain signal is not corrected.

In summary, I demonstrated alterations in networks characteristics resulting from not correcting for global signals. Such alterations include increased connections along the interhemispheric fissure and isolated nodes and subgraphs around the white-matter tracts. However, incomplete global signal correction or lack thereof may not alter some brain network modules, such as DMN. Thus, each practitioner of brain network analysis, especially dealing with networks in voxel-level, should consider the results presented in this work and select an appropriate correction method that is suitable for his / her study.

## 5. Acknowledgements

This study is supported in part by the National Institute of Neurological Disorders and Stroke (NINDS) (NS070917). I would like to thank the three reviewers for their insightful comments in revising this paper. Additional details on MRI data acquisition have been added because of the reviewers’ suggestions. Analyses on nodes adjacent to white matter tracts have been added also because of the reviewers’ suggestions. The names of these insightful reviewers can be found on the first page of this article.

